# Development of a bispecific Antibody–Drug Conjugate targeting CD7 and CD33 to treat Acute Myeloid Leukaemia

**DOI:** 10.1101/2022.07.17.500350

**Authors:** Andrew D. Duckworth, Cath Eberlein, Victoria Pollard, Richard Bethell, Oliver Schon, Joseph R. Slupsky, John F. Woolley, Tiffany Thorn (nee Daniels)

## Abstract

Acute myeloid leukemia (AML) is a heterogeneous malignancy of the bone marrow associated with poor outcomes and limited treatment options available to patients. Recent developments have demonstrated that patient stratification based on disease classification (e.g. somatic mutations) allows for selective treatment regimens and greatly improved outcomes. AML patients can also be stratified based upon the heterogeneous immunophenotype of surface antigen expression on leukemic blasts and stem cells. Here we present data identifying a sub-population of AML patients showing expression of both CD33 and CD7 on their tumour cells, using mass cytometry. This combination of antigens is disease-specific, and is not expressed on healthy haematopoietic cells in patients. We developed a bispecific antibody–drug conjugate (ADC) targeting both CD7 and CD33 and demonstrate that these bispecific ADCs are cytotoxic to AML cells *in vitro.* Importantly, the anti-CD33/CD7 bispecific ADCs are more selective than single-antigen targeting ADCs and safely discriminate tumour from healthy cells (either myeloid or lymphoid). These anti-CD33/CD7 bispecific ADCs are well tolerated and selectively target AML cells in pre-clinical *in vivo* studies in mice. This study presents the first proof-of-principle for targeting specific and unique combinations of surface antigens on tumour cells with potential to overcome observed toxicities with current ADCs.

## Introduction

Acute myeloid leukaemia (AML) is an aggressive malignancy of the bone marrow, characterized by uncontrolled proliferation of undifferentiated myeloid lineage cells. It is the most common acute leukaemia and has dismal survival rates in adult patients (Siegel *et al*., 2019; Shah *et al*., 2013). Treatment regimens for this disease typically consist of intensive remission induction chemotherapy (cytarabine and anthracycline). Recently, novel small molecule inhibitors have been developed to tailor treatment for subpopulations of patients who carry specific contributing mutations. Such targeted therapies include midostaurin for patients with a *FLT3* mutation, or ivosidenib and enasidenib for patients who carry either IDH1 or IDH2 mutations, respectively (Stone *et al*., 2017; DiNardo *et al*., 2018; Stein *et al*., 2017). While initial response rates to treatment are high, almost all AML patients who initially achieved a complete response (CR) will relapse.

Recent advances in single-cell sequencing have allowed researchers to understand the mutational heterogeneity in AML. It is clear from these studies that the malignant clone in this disease is, in fact, highly heterogeneous, where evolving sub-clonality is brought about by treatment (Miles *et al*., 2020; Morita *et al*., 2020). Nevertheless, it has been understood for decades that the immunophenotype of AML is complex, dynamic and characterized by interpatient heterogeneity (Costa *et al*., 2017). AML is organized in a cellular hierarchy; the bulk of malignant, undifferentiated myeloid lineage cells that define AML arise from, and are sustained by, a rare population of progenitor cells called leukemic stem cells (LSC). LSCs inAML represent a low-frequency subpopulation of leukemia cells that possess stem cell properties distinct from the bulk leukemia cells including self-renewal capacity, long-term clonal propagation, differentiation potential, and quiescence (Lapidot *et al*., 1994). Thus, a successful and remission-free treatment of AML will only be achieved with therapies that eradicate the bulk of the disease, including LSCs (Pollyea & Jordan, 2017). Novel combinations of cell surface-expressed antigens distinguishing bulk AML cells and LSCs from their healthy counterparts could provide a unique and disease specific *‘finger print’* of malignant cells. This concept has the potential to eradicate the disease without damaging normal haematopoiesis and consequently will benefit the patient.

Antibody-drug conjugates (ADCs) consist of a monoclonal antibody (mAb) targeting a cell-surface antigen conjugated to a cytotoxic payload by a stable linker chemistry. Upon binding to the target antigen and internalisation of the antibody, the payload is released and kills the cell. This strategy has two clear advantages over conventional, systemic chemotherapy: (i) malignant cells are being targeted specifically within the complex mosaic of healthy and diseased cells and tissues and thus, (ii) enables the use of highly and ultra-potent cytotoxic agents that would otherwise not be tolerated by the patient in conventional chemotherapy programmes. To date, an increasing number of ADCs have been approved by the regulatory authorities worldwide for the treatment of cancer (Drago *et al*., 2021). Gemtuzumab ozogamicin (GO; Mylotarg) was the first FDA-approved ADC to treat AML, and targets CD33-expressing AML cells with the cytotoxic payload calicheamicin (Amadori *et al*., 2016; Bross *et al*., 2001). GO has yielded strong clinical efficacy (Hills *et al*., 2014), but is associated with serious adverse effects such as persistent thrombocytopenia, veno-occlusive disease/sinusoidal obstruction syndrome (VOD/SOS), infusion-related reaction, tumor lysis syndrome (TLS) and myelosuppression (Cortes *et al*., 2020; Lambert *et al*., 2019). Indeed, severe myelosuppression has been described in older AML patients treated with low-dose GO as a single agent (Amadori *et al*., 2016). Thus, there is a clinical need to reduce toxicity in those otherwise underserved patients by increasing the specificity of ADCs to cancer cells in AML.

Here, we describe the *in vitro* and *in vivo* efficacy of a first-in-class bispecific ADC selectively targeting AML malignant blast cells that co-express a novel combination of cell surface antigens, namely CD7 and CD33. CD33 is expressed by the majority of myeloid cells whereas CD7, a lymphocytic cell marker, is aberrantly expressed on AML blasts and LSCs (Haubner *et al*., 2019). Our data introduce bispecific ADCs as an improved therapeutic treatment option for AML cancer patients in particular, but as a concept that is applicable to potentially hard-to-treat both blood and solid cancer subtypes more generally.

## Results

### Mass cytometry identification of CD33^+^/CD7^+^ populations of malignant cells in AML patient samples

We examined the immunophenotype of bone marrow samples from 25 newly diagnosed AML patients by mass cytometry using a panel of 38 antibodies that could profile the surface antigen phenotypes within both bone marrow and peripheral blood cells. Leukaemia samples were randomly assigned into 1 of 4 batches, with each batch being pooled together using TeMal live-cell barcoding (Willis *et al*., 2018). To control for batch variation, a repeat healthy peripheral blood mononuclear cell (PBMC) sample was included as a technical reference. After event clean-up and live-cell identification, viable AML bone marrow cells (including cells from the 4 PBMC batch controls) were clustered using FlowSOM into 30 metaclusters (MC) which identified populations of cells that share similar antigen phenotypes (Figure 1A). MCs (i.e. rows) are ordered by their median antigen expression by unbiased hierarchical clustering and annotated according to cell phenotype. The MCs were broadly grouped into progenitor myeloid (annotated in green), myeloid (black) and non-myeloid cell types which were each given their own colour coding. The second heatmap (right side) shows the corresponding abundance of each of these MCs within each colour-coded sample (columns). Samples with similar antigen phenotype abundances were grouped together using hierarchical clustering, with all healthy PBMC samples reassuringly clustering together with low Euclidean distance. Predicted properties of each MC are highlighted in the 2 checkerboards between the heatmaps: the first coloured MCs orange or green according to whether they were associated with a malignant or healthy cell type; the second checkerboard was annotated in black, grey or white, respectively, if the MC was predicted a strong, weak or not a target for an anti-CD33/CD7 dual-specific antibody.

**Figure 1.**
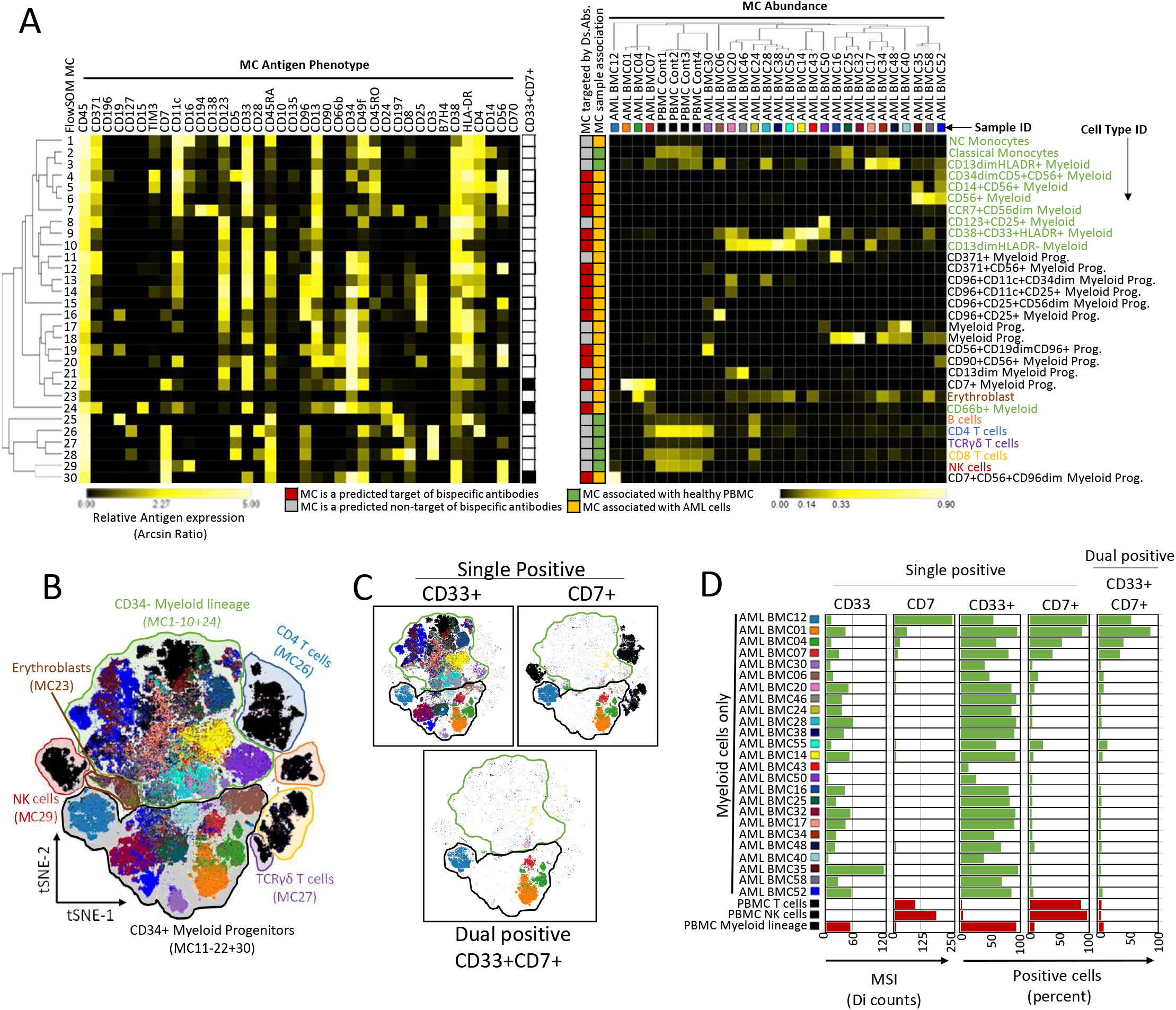
Multivariate analysis of AML BMC samples identifies myeloid phenotypic subtypes with abnormal dual antigen expression. Singlet cells from 25 AML BMC patient samples and 4 replicates of a healthy PBMC sample (used as a batch control) were clustered using FlowSOM into 30 metaclusters (MC) and visualised in 2 dimensions using viSNE. (A) Heatmaps illustrating MC phenotypes (left-hand side) and their relative abundance in each sample (right-hand side). Sample IDs are colour-coded as indicated. In-between each heatmaps, the MCs are annotated for their: i) predicted binding of each proposed dualspecific antibodies (Ds.Abs.; black and white squares); ii) overall potential to be targeted by at least one Ds.Abs. (red and grey squares), and iii) association with each sample type (orange and green squares). MCs are annotated and colour-coded as cell types along the right-hand side, where green text indicates a CD34^-^ myeloid phenotype; black text annotates a CD34^+^ myeloid progenitor phenotype. (B) Dot plot of viSNE analysis, where each dot represents a cell which has been coloured according to its sample ID (as in (A)). Location of cell types identified in (A) are colour-coded and labelled. (C) viSNE plot as in (B) but only events which have the indicated single or dual positivity are shown. Manually gated thresholds for antigen positivity were set using known negative control cell types within the healthy PBMC sample. The CD34^-^ and CD34^+^ myeloid gates are drawn for reference. (D) Bar charts showing the mean signal intensity and percent single/dual positive cells detected within the myeloid populations of each AML sample (row, coloured green). Red bar charts show mean signal intensity and percent single/dual positive cells of the indicated reference cell types present within the healthy PBMC control sample.

To further aid visualisation of this multidimensional dataset, we performed dimension reduction analysis using viSNE and coloured each event within the tSNE dot plot according to the MC from which they were derived (Figure 1B). In addition, to help orientate the reader, we overlay the dot plot with the broad cell type classifications identified from the MCs (these classifications are highlighted using the cell type colour scheme from Figure 1A). This projection was then used to reveal gated populations of cells that were single (top row) or dual (bottom row) positive for CD33 and CD7 expression (Figure 1C), as determined by batch-specific manual gating of CD33 vs. CD7 plots (Supplementary Figure S1). Importantly, CD33 is expressed in multiple MCs across CD34^+^ and CD34^-^ cells, whereas the added combination of CD7 is restricted in this cohort of samples to CD34^+^ AML progenitor cells only. Whilst the majority of AML myeloid cells expressed CD33 (Figure 1D; 10-97%), we were able to confirm CD33 expression levels on this cell population to be similar or marginally lower when compared to myeloid cells in PBMC samples from healthy donors. In contrast, CD7 expression was mostly negative or very low with the notable exception of samples AML-BMC12, 01, 04 and 07; these patient samples showed large percentages of myeloid cells that were positive for this marker (35-90%-range of myeloid cell population). Crucially, within these same samples, distinct populations of dual positive cells could clearly be identified. Taken together, these data show that by using mass cytometry we can i) identify distinct populations of dual positive myeloid cells in AML patient samples & ii) establish the unique twin antigen pair CD33/CD7 to be present on the same cells in a significant subpopulation of AML patient samples.

### A distinct CD33^+^/CD7^+^ population of cells is absent in bone marrow samples from healthy donors

To determine whether CD33 and CD7 dual antigen expression was present only on AML cells but not on non-malignant cell populations, we profiled 4 bone marrow and 7 PBMC samples from healthy donors for antigen expression. PBMCs were pooled into one sample tube using CD45 live-cell barcoding (Hartmann, *et al.* 2018) prior to immunophenotyping by CyTOF. As before, we used the same panel of antibodies for our AML sample analysis presented in Figure 1A. After data clean-up and dead cell exclusion, data events were dimensionally reduced into two dimensions and clustered using viSNE and FlowSOM, respectively. To avoid undersampling of events, and to maximise identification of rare cell types, PBMCs were split into CD3^+^ and CD3^-^ events to produce 3 total “sample types” prior to multidimensional analysis. Figure 2A, C and E show the viSNE plots for each of these sample types, in which events have been arbitrarily coloured according to their FlowSOM MC followed by manual annotation for cell types according to their respective antigen expression (see also Supplementary Figure S2). Bone marrow samples showed distinct cell populations that were not present in PBMC samples. This included CD34^+^ progenitor cells, plasma cells and a significant proportion of non-leukocyte cells. Upon examining bivariate dot plots for CD33 vs CD7 expression, it was immediately evident that CD33^+^/CD7^+^ events, if present, were rare amongst the different healthy sample types. To quantifiably identify specific antibody binding away from the background staining, we applied a bivariate back-gate to identify 1% of total events that were CD33^+^/CD7^+^ (Figure 2B, D, and F). These 1% of events were then visualised by overlaying them back onto each corresponding tSNE analysis (Figure 2B, D, and F) alongside enrichment quantification for each MC cell type (Supplementary Figure S3). In general, the 1% of rare events present across the entire CD33^+^/CD7^+^ gate showed a random spread across all single positive CD33 and CD7 populations on the tSNE map, respectively, suggesting that the majority of these events are due to non-specific antibody binding. Two small subpopulations stood out from this random distribution; about 10-12% of NK2 and 1-5% of pDC present in healthy PBMCs showed CD33^+^/CD7^+^ staining (Figure 2E/F). As previously reported, NK2 cells can exhibit an aberrant transcriptomic, stress-induced profile, especially in AML-derived patient samples (Crinier *et al.*, 2021). The NK2 sub-cell class has been described to play a role in the defence to pathogen insult by cytotoxic activity. In a clinical setting, the potential treatment toxicity profile by removing this subset of NK cells might be viewed as minimal and manageable (Kimura & Nakayama, 2005).

**Figure 2.**
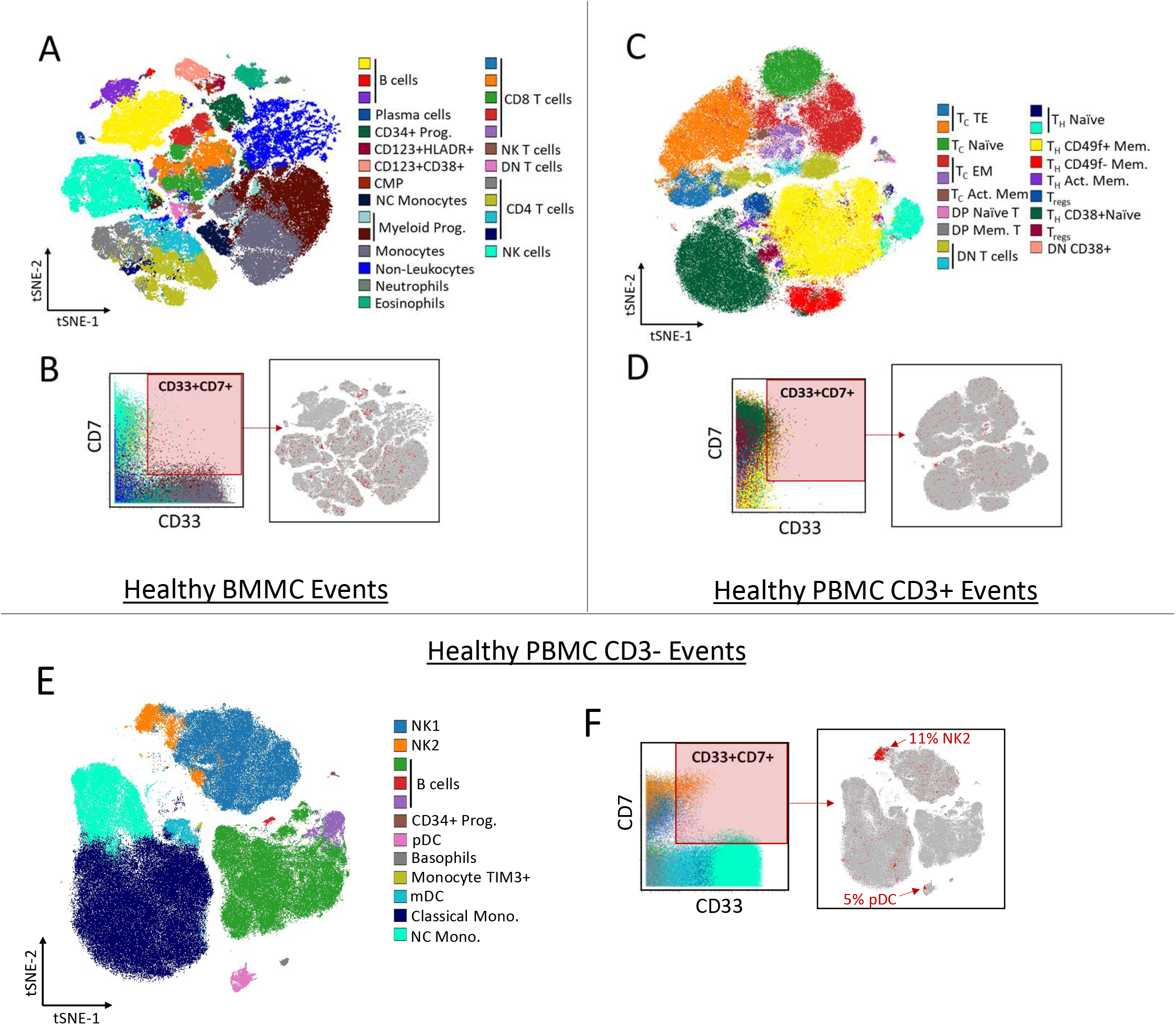
Targeted dual-antigen expression within Healthy BMMC and PBMC. 4 healthy BMMC and 7 healthy PBMC samples were phenotyped for antigen expression using CyTOF. PBMC samples were pre-gated for either CD3^+^ or CD3^-^ events prior to multivariant analysis. (A), (C), and (E) show dot plots of the viSNE landscape for singlet BMMC, CD3^+^ PBMC, and CD3-PBMC, respectively. Each cellular event is coloured for the FlowSOM MC to which it belonged and each MC has been ascribed a descriptive cell type. (B), (D), and (F) show corresponding analysis for (A), (C) and (E), respectively. Bivariate plots and dual expression gates of the antigen combination are shown with events colour-coded for their corresponding sample specific MCs. Dual-antigen positive events are overlayed in red onto their corresponding viSNE plot, while dual negative events are coloured in grey. Cell types that showed significant percentage (>2%) of dual positive cells above non-specific background levels are presented in (F).

Taken together, the data in Figures 1 and 2 show that bone marrow and PBMC tissues from heathy donors do not contain distinct populations of CD33^+^/CD7^+^ cells in the same way as significant subpopulations of patients with AML do. Although there are mature cell populations in healthy donors that are CD33^+^/CD7^+^, these cells are rare and make up only a small fraction of this specific cell type. Our findings therefore present a *twin antigen fingerprint* in AML patients with a unique CD33/CD7 antigen combination present on blast cells but found only very rarely in healthy blood populations. Based on these results, we chose the combination of CD33 and CD7 as a viable twin antigen pair in AML to develop a bispecific antibody drug conjugate with the potential for future clinical application.

### Generation of bispecific antibody fragments targeting CD33 and CD7

In order to generate a bispecific antibody binding fragment (bi-Fab) to target CD33^+^/CD7^+^ cells we used standard click-chemistry, and exploited a framework engineered cysteine as a reactive site, to join recombinant Fab proteins of anti-CD33 and anti-CD7 antibodies. In this format we expected a significant avidity component of antigen binding if the engineered bi-Fab encounters CD33^+^/CD7^+^ cells. To take advantage of this we therefore decided to modulate the individual binding affinity of each Fab entity to its target. We used site-directed mutagenesis of the antigen-binding codons within heavy and light chain coding sequences of the respective Fab proteins to change their respective binding affinity. The anti-CD7 and anti-CD33 Fab variants were expressed in mammalian cell culture, purified and their antigen binding affinities determined by surface plasmon resonance (SPR; Supplementary Figure S4A). This approach yielded 12 mutant anti-CD7 Fabs with affinity reductions in the range of 2-to 152-fold compared to the parental anti-CD7 Fab. For anti-CD33 Fabs, 10 mutants were selected for expression; these were found to have affinity reduction in the range of 2- to 2000-fold compared to the parental Fab (Supplementary Figure S4B). Selected pairs of anti-CD7 and anti-CD33 Fabs, both mutant and wild type, were used to produce a series of affinity modified bi-Fab ADCs (Supplementary Figure S4C). Framework cysteine residues (either cognate or engineered) were used for i) Fab cross-linking and ii) cytotoxic linker/payload conjugation using standard maleimide chemistry.

### BVX100 binds preferentially to cells expressing both, CD33 and CD7

To evaluate the biological effects of different bi-Fab ADC candidates in cells expressing different levels of CD7 and CD33, we sought to identify, characterise and establish cell lines with different antigen expression profiles. We determined the absolute antigen numbers expressed on 15 cells lines that, collectively, represent all combinations of CD7 and CD33 expression patterns and exhibit a broad range of absolute and relative expression levels (Supplementary Figure S5A).

In order to determine whether 2 different binding arms on the bi-Fab allowed preferential binding to CD7^+^/CD33^+^ cells when compared to monovalent anti-CD7 and anti-CD33 Fabs, we conducted cell binding assays. Here we used unconjugated BVX100 and the corresponding parental anti-CD7 and anti-CD33 monovalent Fab proteins from which it had been generated. The bi-Fab and monovalent Fabs bound only to cells that expressed the relevant antigens (Supplementary Figure S5B). As expected, BVX100 bound to a greater extent to dual positive cell lines (CD7^+^ and CD33^+^) compared to monovalent, mono-specific Fabs. In the case of cells expressing only one of the two antigens, the level of binding of the bi-Fabs was similar to that of the corresponding mono-Fab, supporting the notion that affinity and selectivity of individual Fabs had not been compromised in the bispecific molecule format. In cell lines expressing both CD33 and CD7, the relative binding affinity of BVX100 correlates to the respective level of expression (Supplementary Figure S5C). These data suggest that the bi-Fab binds preferentially to cells expressing both antigens due to greater avidity.

We next compared the binding of the heterodimeric anti-CD7/anti-CD33 bi-Fab with corresponding homodimeric bi-Fabs. Here two anti-CD7 Fabs or two anti-CD33 Fabs were conjugated using the same chemistry as described above (Supplementary Figure S5D). Binding of BVX100 was either similar to, or exceeded the level of binding observed with either of the two homodimeric bi-Fabs on CD7^+^/CD33^+^ cells. Importantly, and in line with expectations, the binding of BVX100 to CD7^+^/CD33^-^ cells was reduced compared to the homodimeric anti-CD7 bi-Fab. However, the binding of BVX100 to CD7^-^/CD33^+^ cells was similar to that of the anti-CD33 homodimeric bi-Fab. This suggests a limited influence of the avidity effect on the binding of the homodimeric bi-Fab to CD7^-^/CD33^+^ cells under these conditions, most likely due to the lower expression level of CD33. Expression levels of CD7 generally were higher compared to CD33 levels on cell lines tested here; this was intentional to mimic expression levels as have been reported on AML patient samples. Overall, the results of the binding experiments demonstrate that bispecific proteins targeting CD7 and CD33 preferentially bind to cells expressing both proteins.

### Bi-Fab ADCs targeting CD33^+^ and CD7^+^ cells show cell-specific cytotoxicity

We next armed BVX100 with the cytotoxic drug monomethyl auristatin E (MMAE), and then compared its activity with an MMAE-armed analogue (gemtuzumab-MMAE) of the marketed, CD33-targeting ADC, Gemtuzumab Ozagomycin (GO, Mylotarg), a key competitor ADC. Figure 3A shows that both BVX100-MMAE and gemtuzumab-MMAE were active against CD7^+^/CD33^+^ cell lines, but the latter did not reach the potency of the former which had a significantly lower EC50. Notably, this potency was reversed when gemtuzumab-MMAE and BVX100-MMAE were applied to CD7^-^/CD33^+^ cell lines. In contrast, only BVX100-MMAE was cytotoxic toward CD7^+^/CD33^-^ cell lines, EC_50_ values could not be determined for gemtuzumab-MMAE for these cell lines. We also armed anti-CD7 and anti-CD33 homodimer bi-Fabs with MMAE and tested them in our system. The anti-CD33 homodimer ADC was a potent cytotoxic agent against Kasumi-3 (CD7^+^/CD33^+^) and SHI-1 (CD7^-^/CD33^+^) cell lines (Figure 3A), suggesting that it is not selective for all cells expressing either or both antigens. The lack of sensitivity of some CD33-expressing cell lines to the CD33 homodimer ADC is consistent with data reported for gemtuzumab-MMAE and other CD33-targeting ADCs (Kung Sutherland *et al*., 2013), and is in contrast to BVX100-MMAE, which is potent in all CD7^+^/CD33^+^ cell lines tested. In a similar way the anti-CD7 homodimer ADC was only effective against 2 of the 3 CD7^+^/CD33^-^ cell lines it was tested on, whereas BVX100-MMAE was a cytotoxic agent for all 3. Significantly, the potency of both these ADCs correlated with CD7 expression on the cell lines (Supplementary Figures S5C and S7). Using antigen expression data we estimate the selectivity of BVX100-MMAE for CD7^+^/CD33^+^ compared to CD7^+^/CD33^-^ cells was ~2.8-fold, excluding SET-2 and ALLSIL cell lines with extra-physiologically high levels of CD7. The selectivity of BVX100-MMAE for CD7^+^/CD33^+^ compared to SHI-1 cells and MV4-11 cells (both CD7^-/^CD33^+^) was ~13.7-fold and ~65-fold, respectively. These data suggest that BVX100-MMAE exhibits promising selectivity for target vs non-target cells. They also suggest that affinity reduction of the CD33 arm will not be needed, as the activity against cells that express only CD33 is weak.

**Figure 3.**
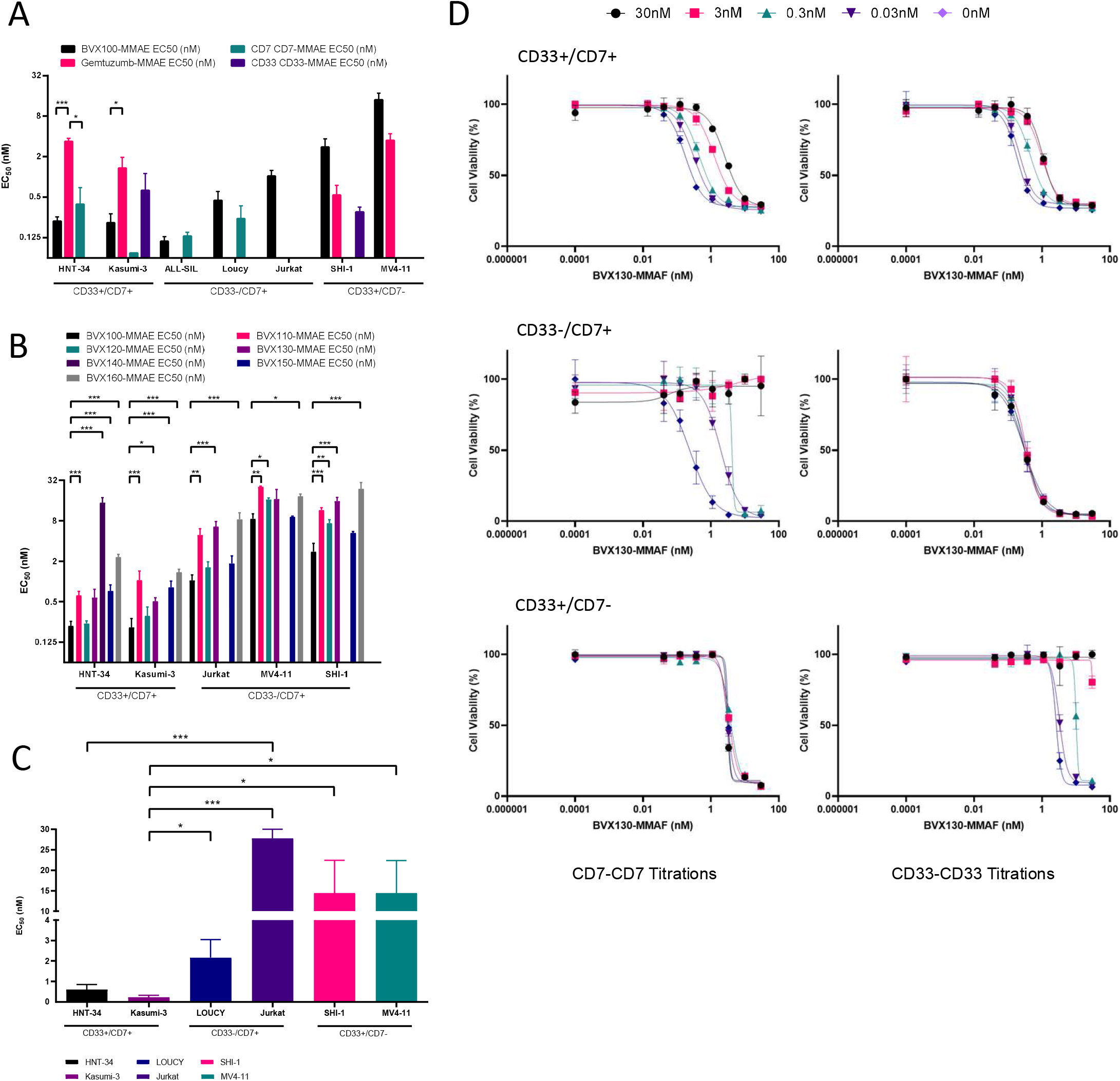
Engineered anti-CD7/anti-CD33 bi-Fab ADCs are selectively cytotoxic against CD33^+^/CD7^+^ cells *in vitro*. (A) Cytotoxicity of BVX100-MMAE, anti-CD7-anti-CD7-MMAE, anti-CD33-anti-CD33-MMAE, and Gemtuzumab-MMAE in leukaemia cell line panels expressing combinations of CD7 and CD33 antigens. (B) Cytotoxicity of a series of novel, affinity modulated bi-Fab ADCs in antigen expressing cell line panels. (C) Selectivity of BVX130-MMAF cytotoxicity for double antigen (CD33+/CD7+) cells. (D) Cytotoxicity of BVX130-MMAF in the presence of increasing concentrations of anti-CD7 or anti-CD33 wild type Fab-homodimers.

Since BVX100 ADC displayed high selectivity for CD7^+^/CD33^+^ cell lines over CD7^-^/CD33^+^ lines, a new series of bi-Fab ADCs was constructed from the mutant Fabs we prepared with the aim of increasing selectivity for double antigen cells further (Supplementary Figures S4 and S6). The majority of the new bi-Fabs contained the original wild type anti-CD33 Fab and 6 different anti-CD7 mutant Fabs which had reductions in affinity in the range 9- to 152-fold. The resulting anti-CD7 mutant panel of bi-Fabs were conjugated to MMAE (Figure 3B) and assessed in cell kill assays (n=3) against a panel of cell lines. Each tested mutant bi-Fab ADC was ranked based on the fold selectivity of CD7^+^/CD33^+^ cell kill over CD7^-^/CD33^+^ and CD7^+^/CD33^-^ cell lines, calculated using EC_50_ values (Supplementary Figure S6). Of the molecules tested, BVX130 had the best overall biological profile with sub-nM potency against HNT-34 and Kasumi-3 cells and substantially higher selectivity for these CD7^+^/CD33^+^ cells compared to Jurkat (CD7^+^; ~13.6 fold), MV4.11 (CD33^+^; ~57 fold), and SHI-1 (CD33^+^; ~41 fold) cells. BVX130 has the affinity reduced anti-CD7 arm and wild type anti-CD33 Fab arm. This bi-Fab was selected for further profiling.

Although the data indicate increased cytotoxicity of BVX130-MMAF (a less soluble payload, requiring active cell endocytosis for cytoxicity, that is more tolerable for *in vivo* studies; Kratschmer & Levy, 2018) in CD7^+^/CD33^+^ double antigen cell lines compared to cell lines expressing either antigen alone (Figure 3C), we wanted to demonstrate that this selectivity required binding of both Fab arms to their respective antigens. To determine this, we carried out cell kill assays using a dose response to BVX130-MMAF in the presence of increasing concentrations of wild type anti-CD7 or anti-CD33 homodimer bi-Fabs which were not linked to a drug (Figure 3D). In CD33^+^/CD7^+^ cells, addition of either homodimer bi-Fab resulted in a decrease in cell kill activity of BVX130-MMAF as indicated by an increased EC50. When CD7^+^/CD33^-^ and CD7^-^/CD33^+^ cell lines were used, BVX130-MMAF was less effective and only respective addition of anti-CD7 or anti-CD33 homodimer BiFabs increased the EC50. Together, these data suggest that both CD33 and CD7 need to be bound by BVX130-MMAF for specific and effective cytotoxicity to occur.

Next, we compared the cytotoxicity of the BVX100 ADC and a gemtuzumab analogue in normal human primary haematopoietic stem cell (HSC)-derived CD33^+^ cells (Figure 4A). Here, BVX100-MMAF did not kill normal CD33^+^ myeloid cells at concentrations up to 10 nM. In contrast, incubation with gemtuzumab-MMAF reduced the number of CD33^+^ cell colonies by 60% when used at 1nM, and by 86% when used at 10nM. Similarly, CD33^+^ monocytes and granulocytes were resistant to BVX130-MMAF when used at concentrations far in excess of its EC50 in CD7^+^/CD33^+^ cancer cell lines (Figure 4B).

**Figure 4.**
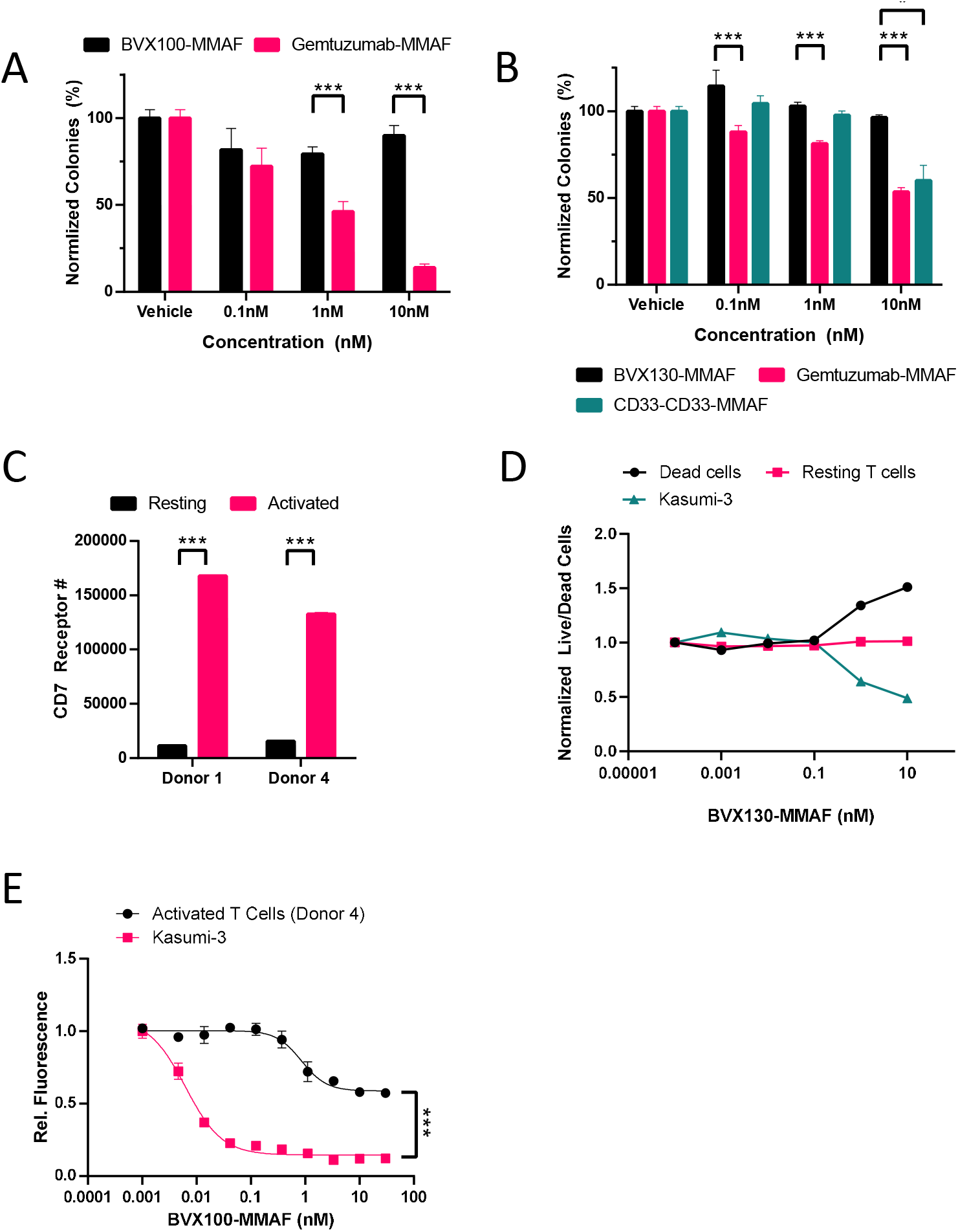
Anti-CD7/anti-CD33 bi-Fab ADCs efficiently target tumour cells while sparing healthy myeloid and lymphoid cells *ex vivo*. (A) Colony forming unit assays of human CD34^+^ bone marrow progenitor cells in the presence of either BVX100-MMAF or Gemtuzumab-MMAF. The bi-valent Gemtuzumab-MMAF kills CD33^+^ cells in a doseresponse manner. BVX test article (monovalent for CD33) does not show colony forming inhibition (B) Colony forming unit assays of human CD34^+^ bone marrow progenitor cells in the presence of either BVX130-MMAF, Gemtuzumab-MMAF or anti-CD33-anti-CD33-MMAF bi-Fab. (C) Quantification of CD7 surface antigen expression on resting and activated PBMC-derived T cells. (D) BVX130-MMAF cytotoxicity in Kasumi-3 (CD33^+^/CD7^+^) cells in mono- and co-cultures of PBMC-derived T cells. (E) BVX100-MMAF cytotoxicity of PBMC-derived T cells compared to Kasumi-3 (CD33^+^/CD7^+^) cells.

The specificity of the bi-Fab-ADCs (BVX100 and BVX130) was further investigated using mixed cultures of Kasumi-3 (CD7^+^/CD33^+^) and normal T cells, comparing resting and activated cells from the latter because of the differential in CD7 expression. To activate normal T cells we incubated them with CD3/CD28 T cell activating beads, and this resulted in 9 to 15-fold increase of CD7 expression levels (Figure 4C). Thus, incubation of BVX130-MMAF with mixed cultures of Kasumi-3 and resting T cells showed that the former were preferentially killed while the latter remained resistant to increasing concentrations of this BiFab ADC (Figure 4D). When we performed the same experiment using mixed cultures of Kasumi-3 and activated T cells we used the more potent, non-affinity reduced BVX100-MMAF. Figure 4E shows that BVX100-MMAF was significantly more cytotoxic towards Kasumi-3 cells than it was towards activated T cells. Taken together with our experiments using normal HSC-derived CD33^+^ cells, these data also suggest that anti-CD7/anti-CD33 bi-Fab ADCs have the potential to selectively kill AML cells at concentrations that would elicit minimal effects on healthy populations and therefore would confer a favourable toxicity profile to patients.

### BVX130-MMAF is stable, well tolerated and cytotoxic to AML cells in vivo

We assessed the safety, tolerability and efficacy of BVX130-MMAF in SCID mice. Figure 5A shows that mice dosed with BVX130-MMAF maintained their body weight over the course of administration, and that this was similar to the response of mice dosed with the standard-of-care AML chemotherapeutic drug cytarabine (Figure 5A). Pharmacokinetic studies of plasma concentrations of BVX130-MMAF showed its presence at levels expected to be cytotoxic to HNT-34 cells (CD7^+^/CD3^+^) was maintained up to 72h following administration of 10mg/kg drug (Figure 5B, Supplementary Figure S8). Of note, clearance of the BiFab-ADC was observed slightly slower at the end of the study than at the beginning. Together, these data confirm that 10mg/kg dosed i.v. twice weekly is a suitable dosing regimen for BVX130-MMAF administration.

**Figure 5.**
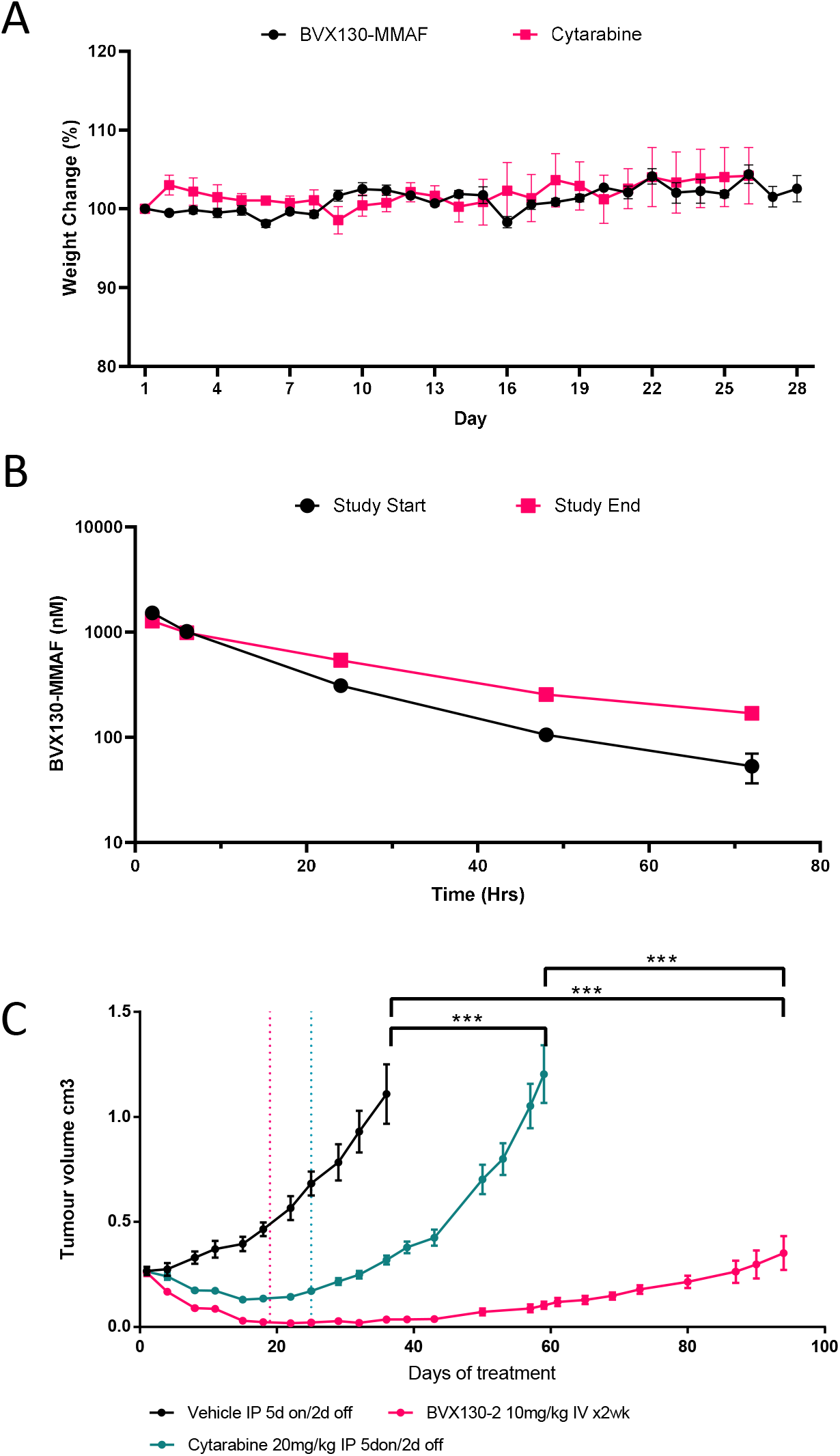
Anti-CD7/anti-CD33 bi-Fab ADCs efficiently target tumour cells *in vivo.* (A) Tolerability as a measure of maintained body weight was assessed in female SCID mice. BVX130-MMAF was dosed at 10 mg/kg twice a week and delivered intravenously for 28 days. For comparison cytarabine was dosed at 20 mg/kg delivered intraperitoneally using a regimin of 5 days on/2 days off for a total 26 days. Body weight was measured daily and normalised to that of the mice on day 0 before administration of drug. (B) Pharmacokinetic data of BVX130-MMAF plasma levels in mice dosed with 10mg/kg drug. Plasma samples were taken at 1h, 24h, 48h and 72h following administration of BVX130-MMAF from mice at the beginning (day 1) and at the end of above tolerability study. Plasma concentration of BVX130-MMAF was measured by ELISA. (C) *In-vivo* efficacy study of BVX130-MMAF in HNT34 (CD33+/CD7+) tumour bearing SCID mice compared to cytarabine and vehicle. 5×10^6^ HNT34 cells were implanted into SCID mice in 50% Matrigel. Each arm contained 10 mice, and tumour volume was assessed at the indicated intervals until termination due to ill health.

Efficacy of BVX130-MMAF was assessed in SCID mice carrying subcutaneous HNT-34 cell tumors (Supplementary Figure S9). Administration of BVX130-MMAF using the above dosing regimen resulted in Tumor Growth Inhibition (TGI) values of 100% (98% regression, p<0.001) and 100% (97% regression, p<0.001) at days 21 and 28, respectively (Figure 5C). Importantly, three animals in this study were assessed as tumor-free at the end of dosing. In comparison, treatment SCID mice carrying subcutaneous HNT-34 cell tumors with cytarabine resulted in TGI values of 100% (47% regression p<0.001) at day 21 (Figure 5C). Consistent with the results of the prior tolerability study, no adverse effects were observed in the BVX130-MMAF treatment group throughout the dosing period. In the cytarabine group, 2/10 animals had body weight losses of >8% on the final day of drug administration, consistent with the data from the tolerability study showing that the dose used in study was the maximum tolerated dose. Re-growth of tumours was monitored after the end of dosing with the cytarabine arm terminated at 57 days due to tumour size. The BVX130-MMAF group was terminated on day 95 as one animal had developed an unrelated lymphoma, and the overall condition of the animals in the group had begun to deteriorate. As has been reported previously (Monticello *et al*., 1994), the SCID mouse model is not stable for extended period of time and spontaneous thymic lymphoma are a consequence of the immune-compromised background of the mouse model rather than an effect of the test article.

## Discussion

Current treatment regimens for AML have essentially remained unchanged for four decades, the combination of standard-dose cytarabine and an anthracycline antibiotic (e.g. daunorubicin) for seven and three days (‘7+3’), respectively. Whilst rates of response to initial chemotherapy in AML are reasonably good, unfavourable karyotypes or specific mutational burden (e.g. TP53 mutations) can push median survival as low as 4 months (Rücker *et al*., 2012; Welch *et al*., 2016). Significantly, older patients have the worst outcomes with 5-year survival rate of <20% (Fernandez, *et al*., 2009; Löwenberg *et al*., 2009). The extreme mutational heterogeneity of AML has hindered the progress to develop alternative therapies. However, in recent years targeted therapies for AML are beginning to become more widespread additions to induction and consolidation regimens. Improvements to AML patient survival has been demonstrated with drugs such as midostaurin and gilteritinib for patients with FLT3 mutations, as well ivosidenib or enasidenib for patients with mutations of IDH1 and IDH2, respectively (Stone *et al*., 2017; DiNardo *et al*., 2018; Stein *et al*., 2017). As our ability advances to profile specific markers of prognosis in individual AML patients more rapidly and more specifically, so will our options for patient treatment. Indeed, it is now clear from the BEAT AML master trial that stratification of AML patients prior to induction chemotherapy, based on molecular profiling, can offer dramatically improved overall survival rates (Burd *et al.*, 2020). This offers a clinically justified route to personalised medicine alongside delayed induction chemotherapy, previously considered unwise for AML patients. Thus, it is likely that improvements over existing therapies for AML will be based on similarly personalised and stratified approaches, tailored to subsets of patients to maximise their clinical benefit.

Here we propose an approach to treatment that bypasses the underlying heterogeneous mutational complexity of AML and instead focuses on the immunophenotype. By utilizing high-throughput mass cytometric profiling of AML bone marrow, we have identified a subset of AML patients whose disease is characterized by a targetable combination of surface antigens, namely CD33 and CD7. We show that a majority of AML patients highly express CD33 and a subset have substantial CD33^+^/CD7^+^ populations (Fig. 1). This combination of CD33 and CD7 antigens presents a therapeutic opportunity, as these antigens are rarely coexpressed on healthy cells (Fig. 2). These data are largely in line with previous studies, as CD33 is typically understood to be a myeloid-expressed antigen while CD7 is expressed on lymphoid cells, only aberrantly expressed on AML blasts and LSCs (Haubner et al., 2019). The expression of CD7 in AML has been associated with unfavourable cytogenetics, reduced leukaemia free survival and relapse, as well as unfavourable clinical outcomes (Abdulateef, *et al.* 2014; García-Dabrio *et al.* 2015; Venditti *et al*., 1998).

Based on these findings we undertook the development of a first-in-class bispecific ADC to selectively target CD7 and CD33. ADCs are a rapidly expanding drug class, with twelve molecules already approved for the treatment of cancers, including Mylotarg for AML (Li *et al*., 2021). All of these approved ADCs target single antigens on tumour cells. In the case of Mylotarg, the ADC targets CD33. In fact about 85–90% of AML cases express the CD33 antigen. However, CD33 is also expressed on healthy, early multi-lineage hematopoietic progenitors, myelomonocytic precursors and more mature myeloid cells (Molica *et al*., 2021). Due to the lack of selectivity of a single antigen-targeting ADC (as decribed earlier) such as Mylotarg, healthy hematopoetic cells are also targeted. For mono-specific treatment regimen, this commonly results in myelosuppression, a severe and dangerous adverse effect (Castaigne *et al*., 2012) frequently observed in clinical settings.

The goal of our study was to overcome this toxicity by designing an ADC that will target a second antigen, increasing the specificity of the ADC for the malignant AML cells. The data we present here in AML cell line models validate our approach; our bispecific ADC format targets CD33^+^/CD7^+^ cells preferentially and specifically over CD33^+^- and CD7^+^-only expressing cells, respectively. This has been shown for both receptor antigens expressed at physiologically relevant levels (Fig. 3). Our proof-of-principle data suggests that bispecificity reduces the likelihood for off-tumour cytotoxicity on normal blood cells *in vivo*. Indeed, our *ex vivo* experiments show no cytotoxic effects of bispecific ADCs on bone marrow derived CD33^+^ cells or on CD7^+^ T cells (Fig. 4).

Our encouraging *in vitro* data were recapitulated using *in vivo* pre-clinical models. We have demonstrated the ability to target CD7^+^/CD33^+^ double positive tumour cells in immunocompromised mice, greatly reducing tumour burden. Here, HNT-34 dual positive cells were specifically chosen for this *in vivo* study because of their low levels of CD33 surface expression. In previous studies (Kung Sutherland *et al*., 2013), this model has been shown to be insensitive to treatment with anti-CD33 monospecific ADCs, such as gemtuzumab ozogamicin, most likely attributable to low CD33 antigen expression in this cell line (Molica *et al.*, 2021). These initial pre-clinical mouse *in vivo* data suggest a superior efficacy in tumour bearing mice compared to current treatment options. Our data also shows that ADC-based tumour targeting is likely a more effective strategy than the standard-of-care chemotherapy for AML, namely cytarabine. Importantly, our studies demonstrate no overt phenotypic toxicity in mice which supports our *in vitro* specificity data.

Taken together, we have demonstrated a novel bispecific ADC capable of effectively, and specifically targeting AML cells in *in vivo* pre-clinical models. The results presented here provide a proof-of-principle for our therapeutic strategy of targeting specific and unique combinations of surface antigens on AML cells with great potential to overcome observed toxicities with current ADCs.

## Methods

### Fab fragment cloning, expression and purification

Fab amino acid sequences containing an engineered free cysteine in the CH1 domain were designed and codon-optimised for expression in mammalian cells (HEK293). Sequences were subcloned into expression vectors provided by external suppliers (Absolute Antibody, Oxford, UK; Peak Proteins, Macclesfield, UK). Transient expression cultures were maintained for between 5-14 days and secreted Fabs purified by cation exchange chromatography followed by dialysis into PBS, pH7.4. QC of purified protein reagents used in subsequent assays met all required parameters of <5% oligomers and purity as assessed by SEC-HPLC & SDS-PAGE, respectively.

### Production of bi-Fabs

In brief, stock Fab proteins were incubated with excess reductant (TCEP). Full reduction and cysteine uncapping was monitored by reverse phase HPLC. The HC/LC interchain disulphide bond was re-oxidised using excess dHAA, leaving the uncapped engineered cysteine in CH1 available for modification. Bi-Fab formation was achieved using well established bio-orthogonal SPAAC chemistry. The chemical reaction was monitored by reverse phase-HPLC under reducing and non-reducing conditions. The bi-Fab product was purified away from unreacted monoFabs and HMW aggregate using preparative size exclusion chromatography with PBS, pH 7.4 as mobile phase. Pooled bi-Fab fractions were concentrated, aliquoted and either used for drug conjugation or stored long term −80°C.

### Linker-Toxin conjugation

Bi-Fabs were incubated with an excess of TCEP (20 eq, Sigma Aldrich, Gillingham, UK) to fully reduce the interchain disulfides, followed by the addition of excess linker-toxin for >2hrs at 35°C. Generation of monovalent Fabs conjugated to linker-toxin was identical apart from capping the reactive engineered cysteine prior to maleimide chemistry. Linker-toxins used in this study are: maleimidocaproyl-val-cit-PAB-MMAE and maleimidocaproyl-MMAF (MedChemExpress, Cambridge, UK). Excess linker-toxin was removed by GF into PBS, pH7. HIC-HPLC (hydrophobic interaction chromatography) was used to monitor both, molecule integrity and number of linker-toxins per protein (Drug-antibody-ratio, DAR).

### BVX130-MMAF in vivo grade

Bulk anti-CD7-6 and anti-CD33 wild-type Fab proteins were treated as above to produce BVX130 bi-Fab stock. BVX130 was then conjugated to mcMMAF as above. Residual endotoxin in the BiFab preparation was removed using an endotoxin removal kit (Pierce: High Capacity Endotoxin Removal Spin Column; Thermo Scientific, UK). Final endotoxin reading was <0.2 EU/mg at a final concentration of 1.02 mg/ml. BiFab-MMAF reagent for tolerability and efficacy studies were aliquoted and stored at −80 °C until use.

### Cell lines and reagents

All cell lines were acquired from DSMZ (DSMZ, Braunschweig, Germany) and cultured in growth media containing RPMI 1640 (Invitrogen, UK) supplemented with 10% heat inactivated Foetal Bovine Serum, (FBS HI; Invitrogen) and 1% Glutamax (Invitrogen). For cell kill assays, 1 % penicillin / Streptomycin solution (Invitrogen) was added. Mouse monoclonal [124-1D1] to CD7 (Abcam, Cambridge, UK); anti-CD33 Monoclonal Antibody (P67.6)-FITC, anti-CD7 Monoclonal Antibody (eBio124-1D1 (124-1D1))-PE, Mouse antiHuman IgG Fab Secondary Antibody-PE and LAMP1 Monoclonal Antibody (LY1C6) eFluor 660 (all from Thermofisher Scientific); CellTiter 96^®^ AQueous One Solution Assay (Promega, Southampton, UK); Human CD34^+^ Progenitor Cells (hCD34+-CB; Promocell, Heidelberg, Germany); Fixation/Permeabilization Solution Kit (BD Biosciences, Wokingham, UK); MethoCult™ H4035 Optimum without EPO (Stemcell Technologies, Cambridge, UK); Cryopreserved Peripheral Blood Mononuclear cells (PBMCs; Cambridge Bioscience, Cambridge, UK).

### Cell binding

Cells were harvested and counted. For each binding condition, 100,000 cells were incubated with primary antibody in PBS / 0.1% BSA for one hour on ice to prevent internalisation. Following incubation, cells were washed in PBS to remove any excess antibody. When required cells were incubated with secondary antibody in PBS/0.1% BSA on ice for 45 minutes. Excess secondary antibody was removed, and cells were analysed for fluorescence using a FACSCalibur (Becton Dickinson, Oxofrd, UK). Data was analysed using Cell Quest software and Geomean plotted.

### Cytotoxicity analysis

Cells were harvested and counted. Cells were dispensed at 20,000 cells per well into 96-well plates or at 2,000 cells per well in 384 well plates. Titrations of MMAE conjugated Fabs were prepared in RPMI, 10% FBS, 1% Glutamax growth media supplemented with 1% penicillin/streptomycin solution. Each titration was prepared at 10x the final assay concentration. Titrations were pipetted into 96-well or 384-well plates using a tenth of the final assay volume. Cell suspensions were prepared and pipetted onto the Fab titrations and plates incubated at 37 °C, 5% CO_2_ for 72 to 96 hours. CellTiter 96 AQueous One Solution (MTS reagent; Promega) was added and the plates incubated for a further 3 hours at 37 °C, 5% CO_2_. Absorbance was measure at 492nm and 690nm on a Spectramax M2 (Molecular Devices, Wokingham, UK) and OD 492-690nm corrected data was plotted using PRISM software.

### Colony Forming Unit (CFU) assay

MethoCult without Erythropoietin (GF H84535; Stemcell Technologies), containing recombinant human stem cell factor (SCF), recombinant human interleukin-3 (IL-3), recombinant human granulocyte colony-stimulating factor and recombinant human granulocyte-macrophage colony stimulating factor (GM-CSF) was used to induce progenitor cell differentiation and proliferation into different myeloid colony types (CFU-GM). Methocult was prepared as described in the manufacturers protocol and aliquots stored at −20 °C. For CFU assays, 1000 CD34+ cells per well were suspended in 1ml of Methocult / IMDM solution containing titrations of BVX100-MMAF, BVX130-MMAF or gemtuzumab-MMAF, respectively. Each treatment was prepared in duplicate and incubated in meniscus-free 6-well plates with the central areas filled with sterile water. Plates were incubated at 37 °C, 5% CO_2_ to maintain constant pH of the Methocult media for the duration of the experiment. Following 10 – 11 days, CFU-GM colonies were identified and counted with the aid of a 6-well grid.

### In vivo efficacy study

All work was carried out under UK Home Office legislation by Alderley Oncology contract research organisation. Female SCID mice (CB17/Prkdc^SCID^) aged 6 weeks old purchased from Charles River, UK were implanted subcutaneously (s.c.) with 0.1 ml of the human acute myeloid leukaemia cell line HNT-34 at 5×10^6^ cells per mouse in RPMI 1640 mixed 1:1 with matrigel. Animals were randomised into groups of 10 once average tumour volume reached approximately 0.2 cm^3^. Animals were dosed intravenously (i.v) twice weekly with BVX130-MMAF formulated in phosphate buffered solution (PBS) for 4 weeks or intraperitoneally (i.p) with either vehicle or cytarabine in 0.9% physiological saline 5 days on, 2 days off for 3 weeks. BVX130-MMAF was supplied as a 1 mg/ml solution as an aliquot per dose by BiVictriX and vials stored at −80 °C until dosing. Cytarabine was prepared fresh weekly and aliquoted into 5 vials which were stored at 4 °C until dosing. All formulations were adjusted at room temperature prior to dosing. Tumours were measured twice weekly by caliper and the volume of tumours calculated. Tumour samples were taken at termination of each group, half the tumour was snap-frozen in liquid nitrogen and stored at −80 °C, whereas the other half was fixed in 10% formalin buffer for 24 hours, transferred to 70% EtOH and then embedded in paraffin.

### Mass Cytometry

All centrifuge steps are performed at 500rcf/5mins/4°C unless stated otherwise. Cryopreserved samples were thawed and mixed with 1mL of 37°C culture media containing RPMI buffer (Sigma Aldrich) containing 10% FCS, L-Glut and penicillin/streptomycin. An additional 8mL of 37°C culture media was then added while agitating and cells were immediately centrifuged at room temperature (RT). Samples were resuspended in 5mL of culture media and then counted using a haemocytometer. 3×10^6^ cells were removed into a separate tube for each sample to be stained for mass cytometry. These samples were washed once in ice cold MaxPAR PBS (Cat# 201058; Fluidigm, London, UK) and then resuspended in the appropriate TeMal barcode (200nM in PBS) or anti-CD45-cadmium barcodes (6 choose 2; 0.5μg/mL in cell staining buffer, CSB; Cat#201068; Fluidigm). Samples were incubated for 10mins at RT with TeMal reagents or 4°C for 30mins with anti-CD45 barcodes. Cells were then washed twice in CSB before barcoded samples were pooled into one tube containing PBS. Then, samples were resuspended at 10^7^ cells/mL in a working solution (1:1000 dilution in RT MaxPAR PBS) of Cell-ID Cisplatin. Cells were left at RT for 5mins, after which 3X volume of CSB was added to each sample before centrifugation. Cell pellets were then resuspended in FcX blocking solution at 50μL/3×10^6^ cells (FcX stock diluted 1:10 dilution in CSB) and incubated at room temperature for 10mins. A 2X concentrated antibody cocktail was directly added to the cells suspended in FcX solution and incubated for a further 30mins, agitating after 15mins. Samples were washed twice in ice cold CSB and once in ice cold PBS before being resuspended in RT 1.6% formaldehyde (1:10 dilution in MaxPAR PBS, Cat# 28906; Thermo Scientific) and incubated at RT for 10mins. Cells were then centrifuged at 800rcf/5mins/4°C and resuspended at 3×10^6^ cells/mL in Intercalator solution (1:2000 dilution of 125nM Cell-ID Intercalator-Ir (Cat# 201192A) in Fix and Perm buffer (Cat# 201067; Fluidigm). Cells were left overnight at 4°C.

The next day, cells were washed once in CSB and twice in CAS solution (Cat# 201240; Fluidigm). Cells were filtered through a 30um filter-top test tube (Cat# 08-771-23; Fisher Scientific), counted, and resuspended at 7.5×10^5^ cells/mL. EQ™ Four Element Calibration Beads were added 1:10 and the sample was analysed using a Helios mass cytometer, collecting 1 million events for each BMMC sample or 3 million events of the pooled PBMC.

Events were normalised against the signal on the EQ™ Four Element Calibration Beads and debarcoded to separate each sample within the TeMal barcode pool. Normalised, individual sample FCS files were then uploaded to Cytobank cloud software for analysis. After sample clean-up (removal of doublets and dead cells) samples were then analysed using FlowSOM and viSNE as described in the Results section.

### Statistical analysis

Data analysis throughout was performed using GraphPad Prism 6.0 for Windows software. Where averages and error bars are indicated these are means and standard error of the mean, or standard deviation where specified. Statistical analyses of pairwise comparisons are by two-tailed, non-paired Students t test and for multiple comparisons by one-way or two-way ANOVA with Tukey’s post hoc multiple comparisons test, as appropriate. Throughout *p < 0.05, **p < 0.01, ***p < 0.001.

---

**Supplementary Figure S1**. Bivariate plots showing CD33 vs CD7 expression in the 25 AML samples and batch control healthy PBMC 3774 sample. Quadrant gates used to identify single and dual positive antigen expression are shown, with gates being batch tailored according to antigen expression on healthy PBMC 3774 CD7+CD33-(T cell) and CD7-CD33+ (monocyte) populations.

**Supplementary Figure S2**. Heatmaps showing antigen expression (columns) within the FlowSOM MCs (rows) from the three analysis sample types: healthy bone marrow (top heatmap); CD3+ PBMC (middle heatmap); and CD3-PBMC (bottom heatmap). On the lefthand side of each heatmp the MCs are given colour coding and cell type annotation, while percent abundance of each MC within each sample is shown on the graph at the righthand side.

**Supplementary Figure S3**. Graphs showing percentage single and dual CD33/CD7 positivity within the three analysis sample types for each biological sample analysed. MC ordering, colour coding, nomenclature and phenotypes are the same as Supplementary Figure S2.

**Supplementary Figure S4**. (A) Antigen binding affinities of mutant CD7 Fabs and fold-reduction in affinity compared with wild type CD7 Fab, determined by SPR. (B) Antigen binding affinities of mutant CD33 Fabs and fold-reduction in affinity compared with wild type CD33 Fab, determined by SPR (C) Summary of mutant bi-Fab ADC’s with associated binding affinity of the individual Fab arms, determined by SPR. (D) Cytotoxicity data for affinity-modified ADCs targeting CD33 and CD7 across a panel of cell lines expression all combinations of these antigens

**Supplementary Figure S5**. (A) Quantitative determination of cell surface antigens on an AML cell line panel by flow cytometry. (B) Binding of wild type (WT) bi-Fab (BVX100) and WT CD7 and CD33 monovalent Fabs to AML cell lines. (C) Correlation of BVX100 binding affinity to the expression of both CD7 and CD33. (D) Binding of wild type (WT) bi-Fab (BVX100) and WT CD7 and CD33 homodimer bi-Fabs to AML cell lines.

**Supplementary Figure S6**. Cytotoxicity data for affinity-modified ADCs targeting CD33 and CD7 across a panel of cell lines expression all combinations of these antigens

**Supplementary Figure S7.** EC50 values of BVX130-MMAF in the presence of increasing concentrations of CD7 or CD33 wild type homodimers. Cells were incubated at 37 oC, 5% CO_2_ for 96 hours with quadruplet repeats of each concentration. Following incubation, 5μl of Alamar Blue reagent was added per well and the plates read following a further incubation at 37 °C, 5% CO_2_ for 4 hours.

**Supplementary Figure S8**. ELISA analysis of PK samples from study AO6819-BVX130-MMAF tolerability. Samples were collected at time points 30 minutes, 2, 4, 8, and 24 hours after first IV dose and a 72-hour sample taken (terminal bleed). All BVX130-MMAF samples were analysed in a CD33 capture ELISA assay. A 10-point dose response of BVX130-MMAF was prepared with a top concentration of 5nM and 1 in 2 dilutions to generate a standard curve for interpolation of sample concentrations at each time point. Error bars represent the SEM across 3 female SCID mice per time point.

**Supplementary Figure S9**. HNT-34 Growth Curves in 5 female SCID mice. 5×106 cells were implanted in 50% Matrigel and grown for 60 days.

## Aknowledgments

The authors would like to thank: George Orphanide as a strategy consultant (co-founder Alderley Park Oncology Development Programme, Alderley Edge, Cheshire SK10 4TG, United Kingdom); Theonie Georgiou for ADC reagent generation (former employee of BiVictriX); Sygnature Discovery fo conducting *in vivo* efficacy model (Sygnature Discovery, Alderley Park, Alderley Edge, Cheshire SK10 4TG, United Kingdom.)

JFW and JRS are funded in part by North West Cancer Research.

## Author Contributions

ADD, CE, VP, JFW all were involved in data generation and/or data analysis. ADD, CE, RB, JRS, JFW, TT were involved in study design.JFW wrote the manuscript. ADD, JFW, JRS, OS, TT were all involved in reviewing and editing the final manuscript.

