## Supplementary material for "Development of a bispecific Antibody–Drug Conjugate targeting CD7 and CD33 to treat Acute Myeloid Leukaemia": Supplemtary Data

### Figure S1

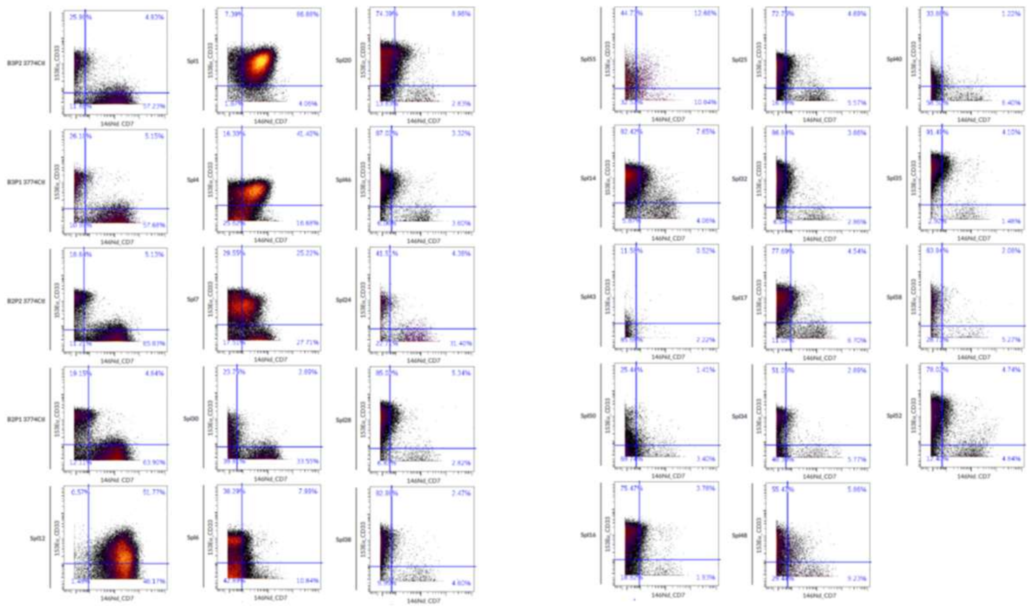

### Figure S2

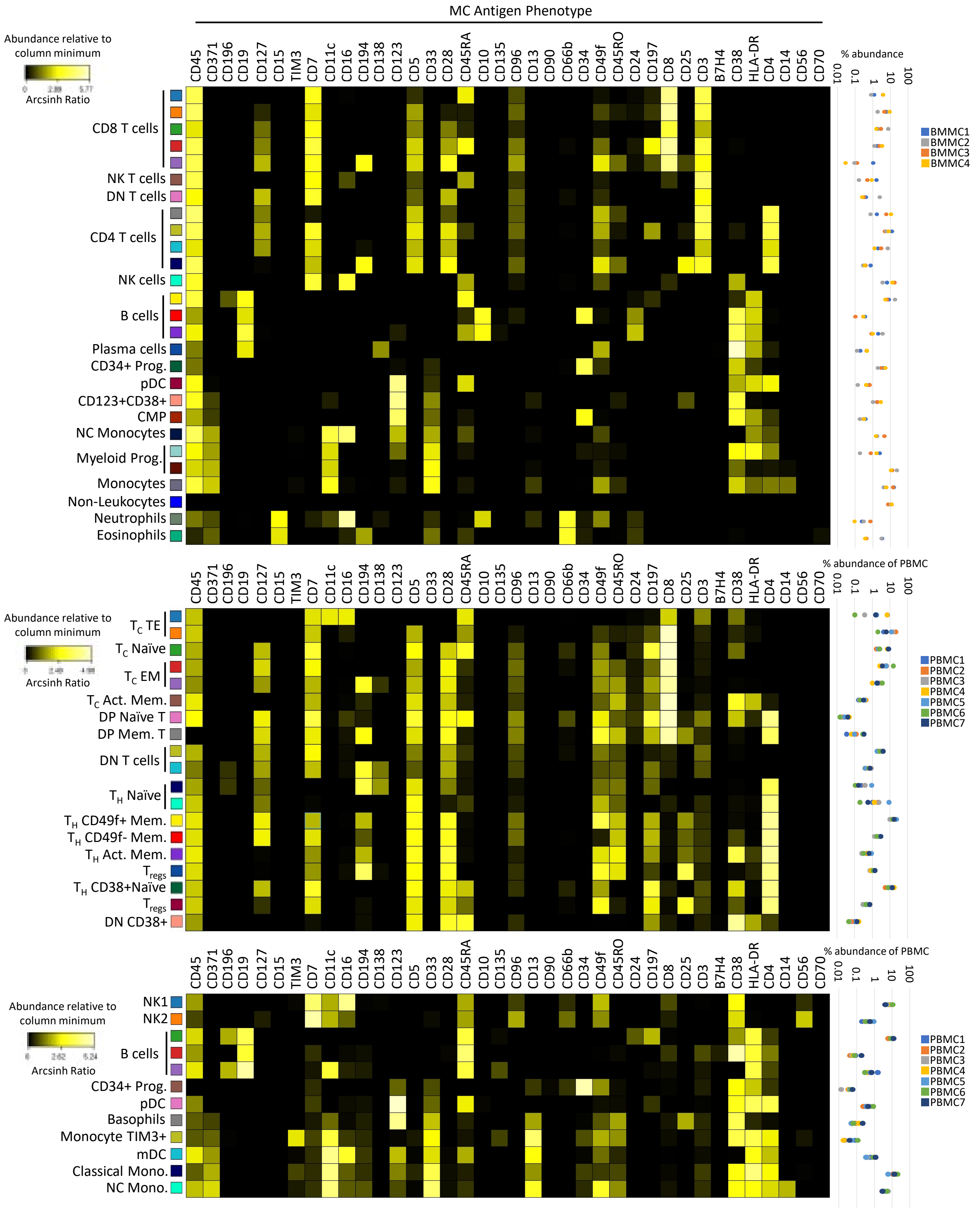

MC Antigen Phenotype
Abundance relative to column minimum
Arcsinh Ratio
CD45 CD371 CD196 CD19 CD127 CD15 TIM3 CD7 CD11c CD16 CD194 CD138 CD123 CD5 CD33 CD28 CD45RA CD10 CD135 CD96 CD13 CD90 CD66b CD34 CD49f CD45RO CD24 CD197 CD8 CD25 CD3 B7H4 CD38 HLA-DR CD4 CD14 CD56 CD70
% abundance
0.01 0.1 1 10 100
CD8 T cells
NK T cells
DN T cells
CD4 T cells
NK cells
B cells
Plasma cells
CD34+ Prog.
pDC
CD123+CD38+
CMP
NC Monocytes
Myeloid Prog.
Monocytes
Non-Leukocytes
Neutrophils
Eosinophils
BMMC1
BMMC2
BMMC3
BMMC4
% abundance of PBMC
TC TE
TC Naïve
TC EM
TC Act. Mem.
DP Naïve T
DP Mem. T
DN T cells
TH Naïve
TH CD49f+ Mem.
TH CD49f- Mem.
TH Act. Mem.
Tregs
TH CD38+Naïve
Tregs
DN CD38+
PBMC1
PBMC2
PBMC3
PBMC4
PBMC5
PBMC6
PBMC7
NK1
NK2
B cells
CD34+ Prog.
pDC
Basophils
Monocyte TIM3+
mDC
Classical Mono.
NC Mono.

### Figure S3

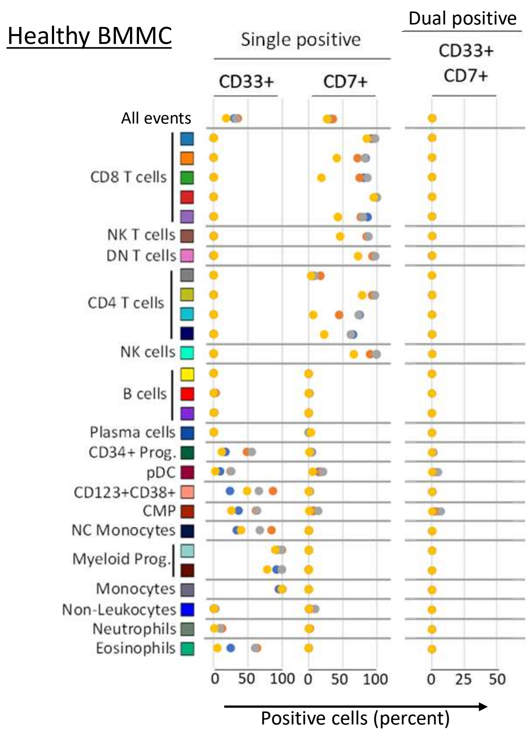
Healthy BMMC
Single positive
Dual positive
CD33+
CD7+
CD33+ CD7+
All events
CD8 T cells
NK T cells
DN T cells
CD4 T cells
NK cells
B cells
Plasma cells
CD34+ Prog.
pDC
CD123+CD38+
CMP
NC Monocytes
Myeloid Prog.
Monocytes
Non-Leukocytes
Neutrophils
Eosinophils
0 50 100 0 50 100 0 25 50
Positive cells (percent)

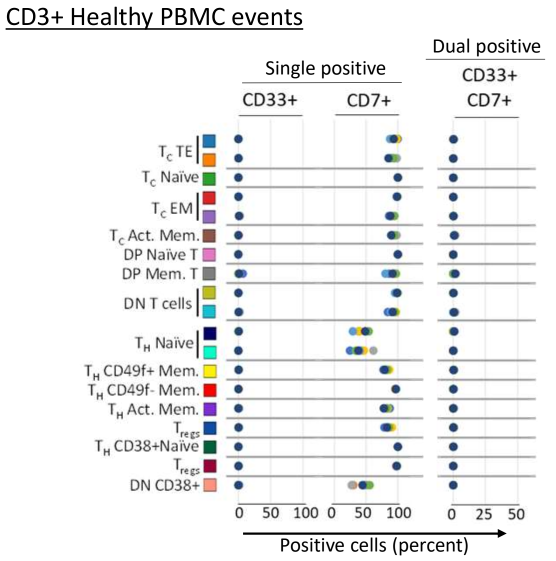
CD3+ Healthy PBMC events
Single positive
Dual positive
CD33+
CD7+
CD33+ CD7+
TC TE
TC Naïve
TC EM
TC Act. Mem.
DP Naïve T
DP Mem. T
DN T cells
TH Naïve
TH CD49f+ Mem.
TH CD49f- Mem.
TH Act. Mem.
Tregs
TH CD38+Naïve
Tregs
DN CD38+
0 50 100 0 50 100 0 25 50
Positive cells (percent)

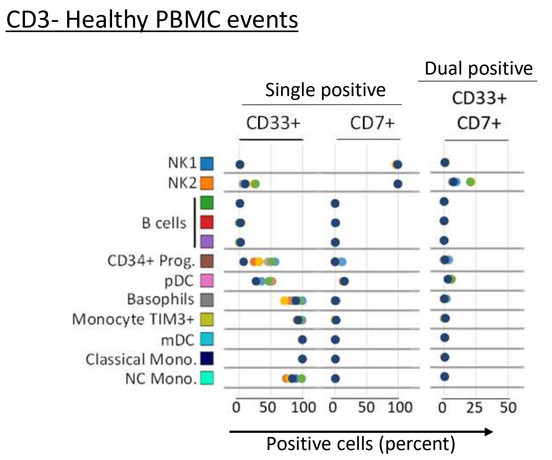
CD3- Healthy PBMC events
Single positive
Dual positive
CD33+
CD7+
CD33+ CD7+
NK1
NK2
B cells
CD34+ Prog.
pDC
Basophils
Monocyte TIM3+
mDC
Classical Mono.
NC Mono.
0 50 100 0 50 100 0 25 50
Positive cells (percent)

### Figure S4

A

| Immobilised ligand | Analyte Number | Mutation | Affinity (nM) | Fold reduction vs Wild type |
|---|---|---|---|---|
| CD7 | 7-1 | Wild type | 3.56 | 1 |
| CD7 | 7-2 | Heavy Chain: Arg57Gly | 75.7 | 21 |
| CD7 | 7-3 | Heavy Chain: Arg57Lys | 32.5 | 9 |
| CD7 | 7-4 | Heavy Chain Tyr102Thr | - | - |
| CD7 | 7-5 | Heavy Chain: Leu105Gly | - | - |
| CD7 | 7-6 | Heavy Chain: Asp104Ala | 45.2 | 13 |
| CD7 | 7-7 | Heavy Chain: Arg57Gly/Tyr108Ala | 541 | 152 |
| CD7 | 7-8 | Heavy Chain: Arg57Lys/Tyr108Ala | 229 | 64 |
| CD7 | 7-9 | Heavy Chain: Tyr102Thr/Tyr108Ala | - | - |
| CD7 | 7-10 | Heavy Chain: Leu105Gly/Tyr108Ala | - | - |
| CD7 | 7-11 | Heavy Chain: Asp104Ala/Tyr108Ala | 246 | 69 |
| CD7 | 7-12 | Light Chain: Gln27Ala | 5.46 | 2 |
| CD7 | 7-13 | Heavy Chain: CDR3 Tyr108Ala | 72.2 | 20 |

B

| Immobilised ligand | Analyte Number | Mutation | Affinity (nM) | Fold reduction vs Wild type |
|---|---|---|---|---|
| CD33 | 33-1 | Wild type | 1.36 | 1 |
| CD33 | 33-2 | Light Chain: Arg35Ser | 1.28 | 1 |
| CD33 | 33-3 | Heavy Chain: Asn100Ala | 1.95 | 1 |
| CD33 | 33-4 | Heavy Chain: Trp102Tyr | 52.7 | 39 |
| CD33 | 33-5 | Heavy Chain: Tyr105Ala / Light Chain: Arg35Ser | 1.75 | 1 |
| CD33 | 33-6 | Heavy Chain: Asn100Ala/Tyr105Ala | 1.72 | 1 |
| CD33 | 33-7 | Heavy Chain: Trp102Tyr/Tyr105Ala | 92.3 | 68 |
| CD33 | 33-8 | Light Chain: Tyr32Asp | 201 | 148 |
| CD33 | 33-9 | Heavy Chain: Asn100Ala / Light Chain: Tyr32Asp | 307 | 226 |
| CD33 | 33-10 | Heavy Chain: Trp102Tyr / Light Chain: Tyr32Asp | 2720 | 2000 |
| CD33 | 33-11 | Light: Chain CDR3 Lys96Ala | 2.23 | 2 |

C

| Bi-Fab Code | Fab arm (Fold affinity reduction vs wild type) | |
|---|---|---|
| | CD7 | CD33 |
| BVX100 | 7.1 (1x) | 33.1 (1x) |
| BVX110 | 7.2 (21x) | 33.1 (1x) |
| BVX120 | 7.3 (9x) | 33.1 (1x) |
| BVX130 | 7.6 (13x) | 33.1 (1x) |
| BVX140 | 7.7 (152x) | 33.1 (1x) |
| BVX150 | 7.8 (64x) | 33.1 (1x) |
| BVX160 | 7.13 (20x) | 33.1 (1x) |
| BVX101 | 7.1 (1x) | 33.4 (39x) |
| BVX102 | 7.1 (1x) | 33.7 (68x) |
| BVX152 | 7.8 (64x) | 33.7 (68x) |
| BVX161 | 7.13 (20x) | 33.4 (39x) |
| BVX162 | 7.13 (20x) | 33.7 (68x) |

### Figure S5

A

| CD7/CD33 Immunophenotype | Cell line | Disease Indication | Specific Antibody Binding Capacity representing receptor number/cell | |
|---|---|---|---|---|
| | | | CD7 | CD33 |
| CD7$^{+}$/CD33$^{+}$ | SET-2 | Essential thrombocythemia | 84893 | 79415 |
| | UOC-M1 | Acute myeloid leukaemia | 61096 | 62943 |
| | HNT-34 | Acute myeloid leukaemia | 22248 | 10907 |
| | Kasumi-3 | Acute monocytic leukaemia | 19595 | 98937 |
| | MOLM-16 | Acute monocytic leukaemia | 3097 | 26249 |
| | HEL-92 | Erythroleukaemia | 1962 | 67636 |
| CD7$^{+}$/CD33$^{-}$ | ALL-SIL | T cell acute lymphoblastic leukaemia | 171872 | NP |
| | LOUCY | T cell leukaemia | 37509 | NP |
| | Jurkat | T cell leukaemia | 15521 | NP |
| | KE-37 | T cell leukaemia | 22599 | NP |
| CD7$^{-}$/CD33$^{+}$ | SHI-1 | Acute monocytic leukaemia | NP | 45597 |
| | MV4-11 | Acute monocytic leukaemia | NP | 230746 |
| | THP-1 | Acute monocytic leukaemia | NP | 10073 |
| CD7-/CD33$^{-}$ | Ramos | Burkitt lymphoma | NP | NP |
| | DND-39 | American-type Burkitt lymphoma | NP | NP |

C

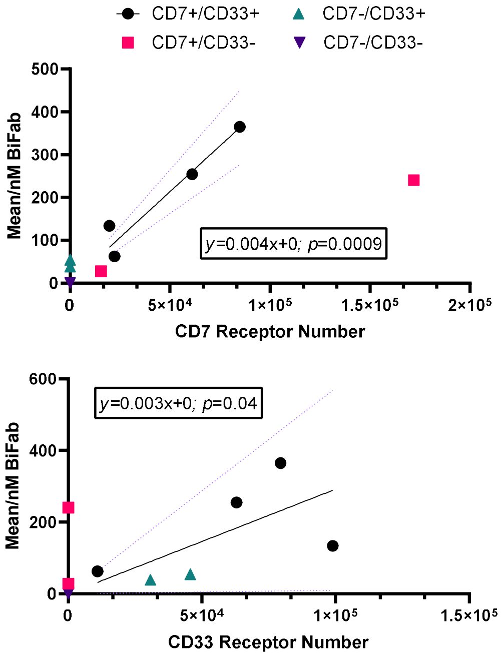


B

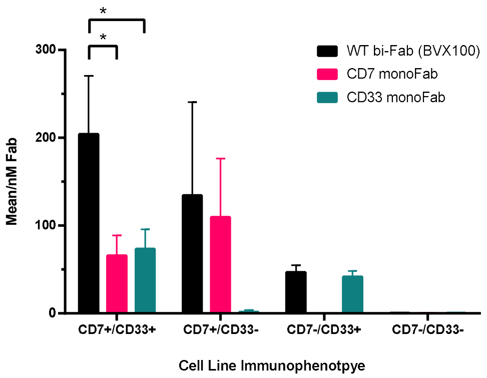


D

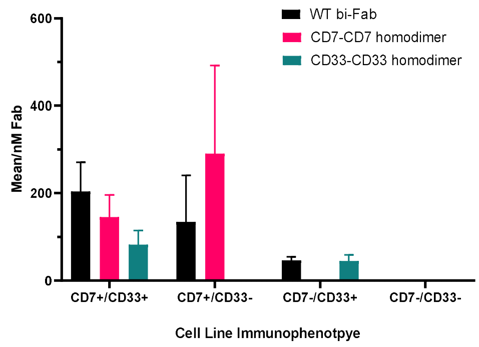

### Figure S6

| Immobilised ligand | EC50 (nM) $CD7^+/CD33^+$ | | EC50 (nM) $CD7^+/CD33^-$ | EC50 (nM) $CD7^-/CD33^+$ | | EC50 (nM) $CD7^-/CD33^-$ | |
|---|---|---|---|---|---|---|---|
| | HNT-34 | Kasumi-3 | Jurkat | MV4.11 | SHI-1 | Ramos | DND-39 |
| BVX100 | 0.19 (±0.04) | 0.14 (±0.07) | 0.92 (±0.22) | 10.77 (±3.69) | 2.26 (±0.89) | NR | NR |
| BVX110 | 0.61 (±0.01) | 0.82 (±0.38) | 4.52 (±1.16) | 26.97 (±1.51) | 11.39 (±0.99) | NR | NR |
| BVX120 | 0.23 (±0.02) | 0.26 (±0.11) | 1.52 (±0.34) | 16.52 (±0.86) | 7.21 (±0.91) | NR | NR |
| BVX130 | 0.40 (±0.19) | 0.45 (±0.07) | 5.83 (±1.25) | 24.07 (±3.25) | 17.48 (±3.28) | NR | NR |
| BVX140 | 17.23 (±4.28) | 24.91 (±3.94) | 22.80 (±5.00) | NR | NR | NR | NR |
| BVX150 | 0.63 (±0.17) | 0.76(±0.20) | 1.60 (±0.54) | 9.19 (±0.17) | 5.23 (±0.29) | NR | NR |
| BVX160 | 2.22 (±0.24) | 1.32 (±0.14) | 7.31 (±2.14) | NR | NR | NR | NR |

### Figure S7

| CD7/CD33 Immunophenotype | Cell line | Receptor Number | | EC50 (nM) | | | |
|---|---|---|---|---|---|---|---|
| | | CD7 | CD33 | BVX100-MMAE | CD7 CD7-MMAE | CD7 CD7-MMAE | Gemtuzumab -MMAE |
| $CD7^+/CD33^+$ | HNT-34 | 22248 | 10907 | 0.19 | 0.25 | NR | 3.35 |
| | Kasumi-3 | 19595 | 98937 | 0.14 | 0.07 | 0.39 | 0.78 |
| $CD7^+/CD33^-$ | ALL-SIL | 171872 | - | 0.11 | 0.13 | NR | NR |
| | LOUCY | 37509 | - | 0.34 | 0.20 | NR | NR |
| | Jurkat | 18089 | - | 0.92 | NR | NR | NR |
| $CD7^-/CD33^+$ | SHI-1 | - | 45597 | 2.26 | NR | 0.29 | 0.42 |
| | MV4-11 | - | 30746 | 10.77 | NR | NR | 2.91 |

### Figure S8

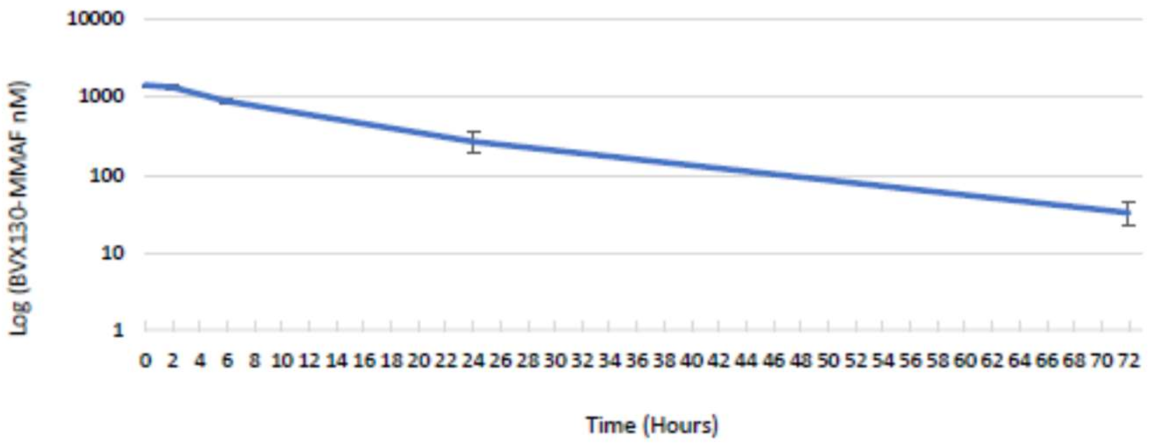

### Figure S9

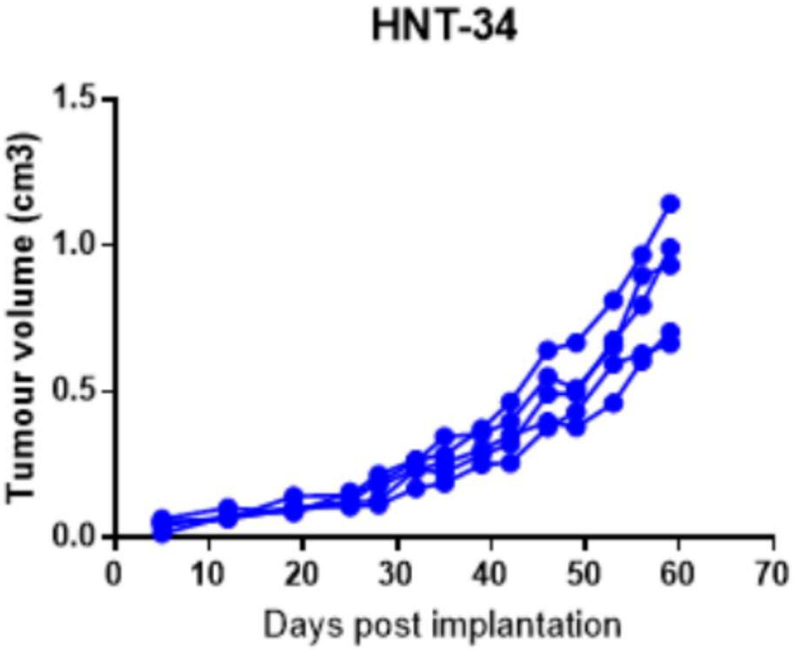
HNT-34
Tumour volume (cm3)
1.5
1.0
0.5
0.0
0
10
20
30
40
50
60
70
Days post implantation
